# From whole-slide histology to ADC maps: Fast diffusion MRI simulation with neural operators

**DOI:** 10.64898/2026.07.22.740029

**Authors:** Iris A. Kohler, Oliver Gödicke, Tristan A. Kuder, Mark E. Ladd, Jürgen Hesser

## Abstract

**Background and Objective:** Simulation of diffusion MRI signals from tissue microstructure is a fundamental problem in quantitative imaging, as it enables controlled study of how cellular architecture influences measured signals. However, physics-based simulations at clinically relevant scales are challenging due to a scale mismatch between imaging and histology: clinical diffusion MRI spans centimeter-scale fields of view with millimeter-scale voxels, whereas histology resolves structure at micrometer scales. Capturing voxel-wise signal formation therefore requires repeated simulations over heterogeneous microstructure, which becomes computationally and memory intensive in classical solvers. We propose a neural operator framework that amortizes this cost by learning local microstruc-ture–signal mappings once and applying them across large tissue regions.

**Methods:** We train a Fourier Neural Operator on finite-element simulations of histology-derived cell segmentations to predict magnetization fields from diffusivity and permeability maps. The model is embedded in a subdomain tiling strategy that enables scalable inference over whole-slide histology images. Unlike most conventional simulation pipelines, inference operates directly on regular grids derived from cell segmentations and does not require meshing.

**Results:** The proposed framework enables simulation of apparent diffusion coefficient maps over 2D liver histology spanning 28.224 mm × 18.144 mm. It achieves over 2,600-fold acceleration compared with CPU-based finite-element simulation, reducing runtime from an estimated 217 days to under 2 hours. The network yields mean relative signal errors of 0.34%–0.43% at high diffusion weighting and 0.03% at low diffusion weighting, with maximum errors below 5%. On manually segmented datasets with greater morphological variability, mean errors increased slightly to 1.37%–1.79%.

**Conclusions:** Neural operators enable computationally practical, mesh-free diffusion MRI simulation by amortizing expensive physics-based computation into a reusable operator applied across local sub-domains. This makes large-scale histology-based diffusion MRI modeling feasible while preserving high accuracy.

## 1. Introduction

Diffusion magnetic resonance imaging (dMRI) provides a noninvasive means to probe tissue microstructure by sensitizing the measured signal to the random motion of water molecules. In practice, dMRI measurements are acquired over multi-voxel imaging volumes, with individual voxels typically spanning up to several cubic millimeters [1]. At these scales, each voxel contains a large and heterogeneous population of cells whose organization determines the measured signal. Physics-based numerical simulations provide a principled framework for studying this microstructure–signal relationship and have been performed using both synthetic substrates and realistic histology-derived geometries [2, 3, 4, 5, 6, 7, 8, 9, 10], employing Monte Carlo [11, 12, 13, 14] and finite-element methods [15, 16] to accurately model diffusion processes within complex cellular environments.

However, dMRI simulations remain computationally demanding, particularly when realistic histology-derived microstructures are considered. A fundamental challenge is the large mismatch in spatial scales between imaging and cell architecture: clinical dMRI spans centimeter-scale fields of view with millimeter-scale voxels, whereas the underlying tissue architecture varies on micrometer scales. Consequently, generating dMRI images requires repeated simulations over a very large number of heterogeneous local microstructures. Considerable effort has therefore been devoted to accelerating large-scale simulations through optimized frameworks such as the Monte Carlo tools RMS [11], MC/DC [12], Disimpy [13], and SpinWalk [14]. These advances have enabled simulations on domains ranging from hundreds of micrometers to millimeter scales, including large cardiac, axonal, and white-matter substrates [17, 12, 18, 19, 7, 20, 21]. Nevertheless, simulating dMRI maps based on large histological regions such as whole-slide images remains computationally impractical when direct numerical simulations must be repeated over many local tissue configurations.

At the same time, machine learning has emerged asa powerful tool for accelerating computational models. In dMRI, most machine learning approaches focus on inverse problems such as reconstruction, signal interpolation, tractography, and microstructural parameter estimation [22]. Comparatively little work has addressed the forward simulation problem itself. One notable example is the polynomial metamodel proposed by [23], which is trained on Bloch–Torrey simulations to predict diffusion tensor–derived scalar metrics from coarse microstructural descriptors. However, this approach bypasses explicit simulation of water diffusion in tissue microstructure by mapping low-dimensional parameter vectors directly to summary dMRI measures, rather than approximating the forward simulation itself.

Recent advances in operator learning provide a promising framework for amortizing expensive numerical simulations. Neural operators learn mappings between function spaces and can approximate solution operators of partial differential equations (PDEs) directly from data [24, 25, 26, 27, 28]. Unlike conventional neural networks operating on fixed-dimensional inputs and outputs, neural operators can generalize across spatial discretizations, geometries, and physical parameters [24, 25]. Among these methods, the Fourier Neural Operator (FNO) [29] has emerged as a particularly effective architecture and has been successfully applied to accelerate simulations in domains including geological storage [30], weather forecasting [31], seismic wave propagation [32], photoacoustic imaging [33], urban microclimate modeling [34], and biomedical engineering [35]. To the best of our knowledge, neural operators have not previously been applied to dMRI simulations.

Although neural operators can substantially accelerate numerical simulations, directly learning dMRI signal formation on clinically relevant domains is itself impractical. Millimeter-scale tissue regions contain thousands of cells and require high-resolution spatial representations, leading to prohibitively large input tensors and correspondingly large memory requirements during training and inference. Moreover, generating sufficiently diverse training data for such domains would require an enormous number of computationally expensive numerical simulations. Consequently, directly learning millimeter-scale dMRI simulations is unlikely to be practical.

A key observation motivating the present work is that dMRI signal formation is governed primarily by local microstructural organization. Within a given tissue type, cellular characteristics such as size distributions, packing density, and compartmental arrangement tend to vary gradually and recur across spatial locations. Consequently, many local tissue regions exhibit similar microstructural patterns and therefore similar relationships between tissue architecture and dMRI signal formation. This recurrence suggests that learning local microstructure–signal mappings may be substantially more efficient than directly learning millimeter-scale simulations.

This idea is closely related to our previously proposed subdomain-based simulation framework [36], which exploited the locality of dMRI signal formation to enable memory-efficient simulations through domain decomposition. However, computational runtime remained dominated by the repeated numerical solution of individual subdomains.

The recurrence of local microstructural patterns suggests an amortized simulation strategy. Rather than repeatedly solving the underlying PDEs independently for each local tissue region, a neural operator can be trained offline on a representative set of microstructures to learn the mapping from local tissue architecture and biophysical parameters to the resulting transverse magnetization field. The computational cost is therefore incurred once during training, after which the learned operator can be evaluated rapidly on a large number of subdomains. This enables efficient simulation over whole-slide images while avoiding repeated numerical solutions.

In this work, we combine a Fourier Neural Operator with the subdomain framework to construct an amortized emulator of local dMRI simulations. The proposed approach learns an operator that maps diffusivity and permeability fields, together with spatial coordinates, to the resulting transverse magnetization field at the end of the diffusion encoding sequence. Combined with a domain tiling strategy, this formulation enables rapid inference directly from histology-derived cell segmentations without requiring mesh generation at inference time. We demonstrate that the proposed framework reproduces finite-element simulations with high accuracy while reducing computation times by more than three orders of magnitude, enabling practical generation of apparent diffusion coefficient (ADC) maps from histological whole-slide images. These results establish a scalable framework for linking histology-derived microstructure to dMRI measurements across large tissue regions.

## 2. Methods

### 2.1. Subdomain framework with neural operator acceleration

This work builds on our previously proposed subdomain framework for large-domain dMRI simulations [36], which enables the computation of dMRI signals over spatial scales far exceeding the memory limits of conventional finite-element solvers. The key observation underlying this framework is that, over the duration of a typical dMRI experiment, diffusing spins explore only a limited spatial neighborhood. The characteristic diffusion length scales as 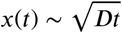, where is the diffusivity and is the diffusionencoding time. As a result, spins contributing to the signal at a given location are insensitive to microstructural features far outside this diffusion length.

Exploiting this locality, the large simulation domain is decomposed into a collection of subdomains. Each subdomain is extended by a fixed margin to form an extended subdomain, which is then simulated independently with no-flux boundary conditions. For sufficiently large extension margins, the transverse magnetization evolution within the central region of the extended subdomain is unaffected by artificial boundaries and matches the behavior that would be observed in a much larger, globally simulated domain. The macroscopic signal is obtained by cropping the predicted magnetization fields to retain only these central regions and aggregating their contributions across the full domain. Specifically, if *M*_*i*_(***x***, *T*_*E*_) denotes the complex transverse magnetization field in the *i*-th extended subdomain evaluated at spatial location ***x*** and echo time *T*_*E*_, and Ω_*i*_ denotes the retained central region corresponding to the original subdomain, the signal is approximated as

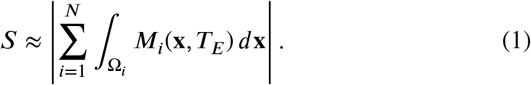

Although the subdomain framework removes the memory constraints associated with large-domain dMRI simulations, the overall computational cost remains high because dMRI simulations must still be performed for a large number of subdomains using conventional numerical solvers. In this work, we replace these explicit simulations with a neural operator that directly predicts the complex-valued transverse magnetization field within a subdomain from its underlying microstructural and biophysical parameters. Figure 1 illustrates the integration of the neural operator into the subdomain framework for generating dMRI signal maps directly from whole-slide histology images.

**Figure 1.**
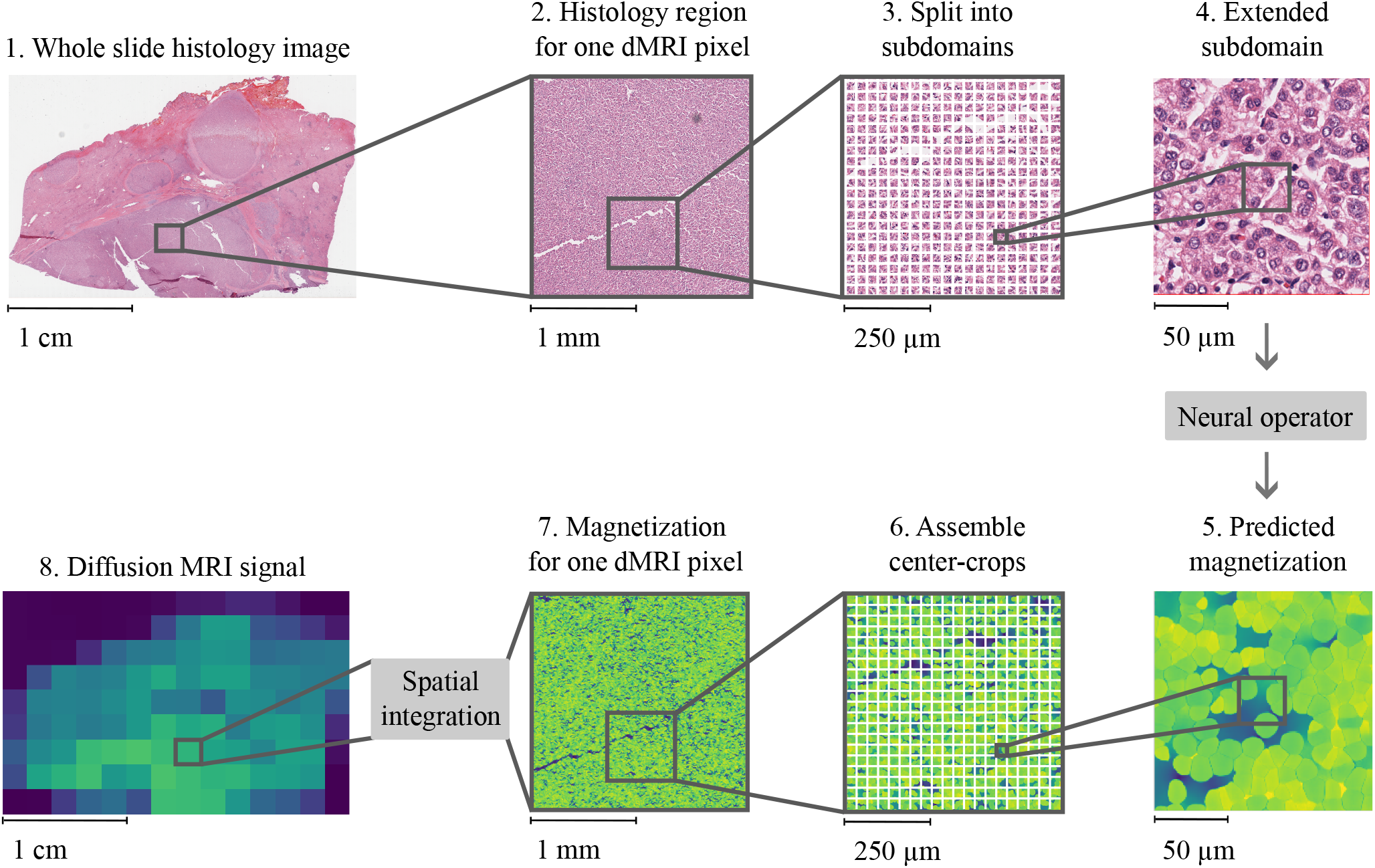
Overview of the proposed framework for large-scale histology-based dMRI simulation. A whole-slide histology image is partitioned into subdomains, each corresponding to a local tissue patch. To reduce boundary effects associated with the finite simulation domain, each subdomain is extended by a fixed margin and provided as input to a neural operator that predicts the complex-valued transverse magnetization field at the end of the diffusion-encoding sequence. For clarity, the figure displays only the magnitude of the predicted magnetization. The full-domain magnetization field is reconstructed by assembling only the central region of each predicted magnetization field, excluding regions potentially affected by boundary artifacts. Spatial integration of the reconstructed field yields the pixel-wise dMRI signal.

The central idea of the proposed framework is that dMRI signal formation is both local and repetitive. Rather than repeatedly solving essentially similar local diffusion problems throughout a whole-slide image, we learn a neural operator that approximates the solution operator once and subsequently reuse it across all subdomains. In this way, the computational burden is shifted from repeated numerical PDE solves to repeated neural network evaluations, allowing practical dMRI simulations across millimeter-scale tissue domains. Although the formulation is general and readily extends to three-dimensional domains, we focus on a twodimensional setting to allow controlled validation against a finite-element reference and direct comparison with our previous subdomain framework.

### 2.2. Training data

To train a neural operator that can be repeatedly applied across heterogeneous whole-slide histology images, we require a dataset spanning a broad range of local tissue architectures. The generation of this dataset comprises three stages: First, selection and segmentation of histological image patches, second, augmentation to increase coverage of the morphology space, and third, finite-element dMRI simulations.

The simulation pipeline itself follows the methodology introduced in our prior subdomain framework work [36]. Briefly, hematoxylin and eosin (H&E)-stained histology images are automatically segmented to obtain cell label images, which are converted into computational meshes and subsequently used for finite-element dMRI simulations with the SpinDoctor toolbox [15]. Only intracellular and extracellular compartments are modeled.

#### 2.2.1. Histology image selection and cell segmentation

We focus on H&E staining primarily due to the widespread availability of whole-slide images (WSIs) covering tissue areas on the order of several square millimeters, which is essential for simulations at spatial scales relevant to clinical dMRI and for our long-term goal of predicting ADC maps from histology.

However, in H&E-stained images, cytoplasm and extracellular space often exhibit similar staining characteristics, leading to ambiguous boundaries during automatic segmentation. In principle, alternative modalities such as fluorescence microscopy or immunohistochemistry can provide clearer contrast between cellular compartments through targeted labeling. In practice, these modalities are typically limited to much smaller fields of view and lack publicly available WSIs, making them unsuitable for large-domain simulations. As a result, H&E represents a pragmatic tradeoff between segmentation fidelity and spatial coverage.

Because the neural operator is intended to be repeatedly applied across heterogeneous tissue microstructure, the training data must adequately cover the range of local cellular configurations expected during inference. We therefore selected 10 WSIs from the TCGA liver hepatocellular carcinoma (LIHC) dataset [37]. From each WSI, tissue patches were sampled on a regular grid with a spacing of 750 µm in both the *x* and *y* directions. This sampling strategy yielded 25 spatial locations per WSI. At each location, multiple segmentation variants were generated (described below), resulting in a total of 10,000 distinct microstructural realizations used for training. The case identifiers and top-left coordinates of the sampling grids for each WSI are listed in Appendix A.

Cell segmentation was performed using QuPath [38] with a pretrained StarDist model [39]. The input histology images were first intensity-normalized by mapping pixel values between the 1st and 99th percentiles. Nuclei were detected at a resolution of 0.5 µm per pixel using a probability threshold of 0.4. Full cell regions were approximated by radially expanding the detected nuclei.

To increase the diversity of cell sizes and packing densities, we generated multiple segmentation variants for each patch by varying the nucleus detection probability threshold (0.4–0.8 in steps of 0.1) and the radial expansion radius (1–15 pixels in steps of 2 pixels). The expansion radius was constrained to be no more than three times the nucleus diameter to prevent overestimation of cytoplasmic extent. The resulting label images were resampled to a spatial resolution of 0.1 µm per pixel to reduce discretization artifacts, particularly along curved cell boundaries.

The dimensions of the extended subdomain and its retained central region used in this work were adopted directly from our previous study [36]. These parameters were determined from reference simulations of free diffusion in a homogeneous domain for the diffusion-encoding protocol used in this work (see Section 2.2.3), in which the magnetization integrated over the retained central region was compared with the analytical free-diffusion attenuation, exp(−*bD*). This analysis ensured that the artificial domain boundaries introduced a negligible error within the retained central region. Specifically, an extended subdomain width of 150 µm and a retained central region width of 32 µm yielded a relative signal error of 0.08% [36].

From the full dataset of 10,000 label images, 1000 samples each were randomly selected and reserved for validation and testing, with the remaining samples used for training.

#### 2.2.2. Dataset composition and training set augmentation

Since the neural operator is expected to operate on previously unseen tissue regions, adequate coverage of the morphology space is essential for robust generalization. Initial experiments indicated limited generalization when training on the original dataset, which exhibited relatively little variation in cellular density and cell size. To quantify morphological variability, we characterized each label image using two features: the cell count and the cell area ratio (fraction of image area occupied by cells). These features are related to histological correlates of dMRI, including cellular density and the intracellular-to-extracellular volume fraction [40].

To increase coverage of this feature space, we applied geometric data augmentation to the 8000 training samples using two strategies: random crop-and-resize operations and crop-and-mirror augmentation, where cropped image regions were mirrored and tiled to reconstruct a full-sized image. Examples are shown in Fig. 2.

**Figure 2.**
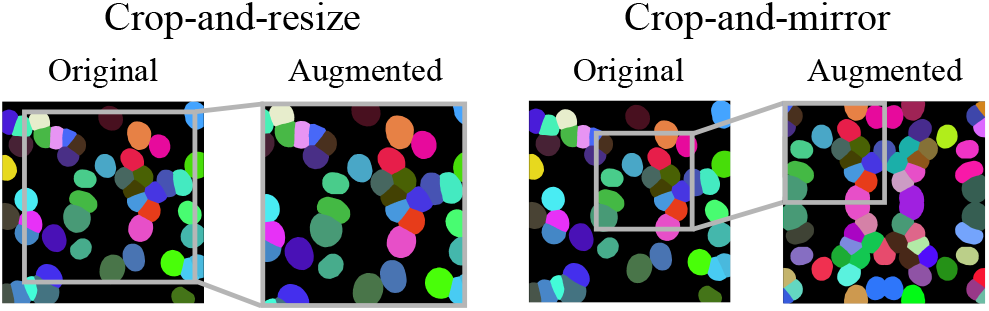
Examples of training data and geometric augmentations. The original segmentation is shown alongside an example of the crop-and-resize augmentation and the crop-and-mirror augmentation.

The original and augmented samples were embedded in the two-dimensional feature space defined by cell count and cell area ratio. To promote a balanced representation of morphological regimes, we binned samples into a 20 × 20 histogram and subsampled within each bin. This procedure yielded a final training set of 4372 images. Figure 3 shows that the augmented and subsampled dataset provides substantially broader coverage of the feature space than the original training data.

**Figure 3.**
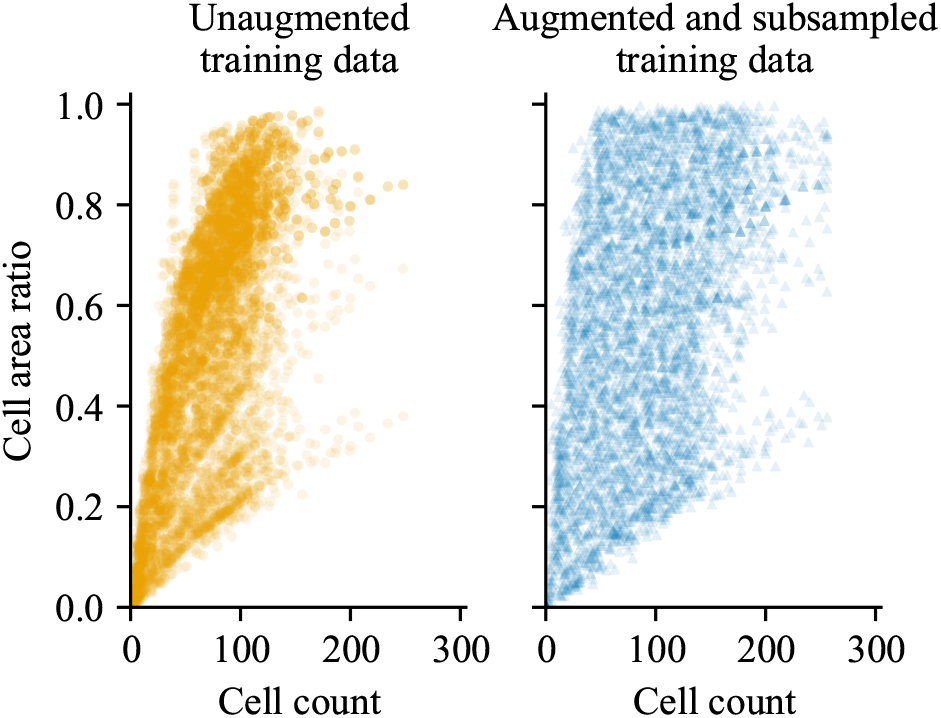
Distribution of cell count and cell area ratio across the unaugmented dataset and the subsampled augmented training set. The subsampled augmented training set shows broader coverage of the feature space than the unaugmented dataset.

#### 2.2.3. Diffusion MRI simulation

We perform the dMRI simulations using the finite element tool SpinDoctor [15], following the numerical setup introduced in our prior study [36]. SpinDoctor requires a triangular mesh which we generate based on the label image with the same procedure described in [36], with a triangle edge length of approximately 0.3 µm. The computational domain is defined on a 150 µm × 150 µm region of interest, and an additional padding of 5 µm is introduced on each side during mesh generation, resulting in a final simulated domain of 160 µm × 160 µm. For the dMRI simulation, we use a pulsed-gradient spin-echo (PGSE) sequence with gradient pulse duration *δ* = 9.5 ms and diffusion time Δ = 25.8 ms. Simulations are performed at diffusion weightings *b* = 50 s/mm^2^ and *b* = 800 s/mm^2^, enabling standard twopoint ADC estimation. The diffusion-encoding gradient is applied along the *x*-axis. The initial spin density was set to 1, and transverse relaxation effects were neglected by setting *T*_2_ = ∞.

For each *b*-value, we generated two complementary datasets to evaluate different aspects of the proposed framework. The fixed-parameter dataset uses a single set of biophysical parameters and is designed to assess whether the neural operator can accurately reproduce dMRI simulations across diverse cellular geometries and spatial organizations. The variable-parameter dataset additionally varies diffusivity and membrane permeability parameters to evaluate whether the neural operator remains accurate across both morphological and biophysical variability.

1. Fixed-parameter dataset: extracellular space diffusivity *D*_ECS_ = 0 003 mm ^2^/s, intracellular space diffusivity *D*_ICS_ = 0.001 mm ^2^/s, and membrane permeability *P* = 0 µm/s.
2. Variable-parameter dataset: extracellular and intracellular space diffusivities sampled uniformly from 0.0002 to 0.003 mm ^2^/s in increments of 1×10^−4^ mm^2^/s; membrane permeability sampled from 0 to 50 µm/s in increments of 2 µm/s.

The selected parameter ranges are consistent with values commonly reported in prior dMRI simulation studies of nonbrain tissues, including liver, lymphocytes, and cardiomyocytes. Specifically, previous works [2, 7, 20, 21, 41, 42, 5, 8] have employed intracellular and extracellular diffusivities spanning 0.0002–0.003 mm^2^ /s, as well as membrane permeabilities ranging from 0 to 50 µm/s. By simulating across this parameter space, we assess whether the proposed neural operator can accurately approximate the transverse magnetization under biophysical conditions representative of those used in the literature.

### 2.3. Neural operator architecture

To eliminate the need for repeatedly solving local finite-element diffusion problems throughout a large tissue domain, we train a neural operator that approximates the mapping from local tissue architecture and biophysical parameters to the resulting transverse magnetization field. For this purpose, we employ the Fourier Neural Operator (FNO) [29], which parameterizes integral kernel operators in the Fourier domain. The architecture follows a standard pattern of 1) a learned lifting transform that projects inputs to a higher-dimensional feature space, 2) a stack of Fourier layers interleaved with nonlinear activations, and 3) a projection back to the target output space. Each Fourier layer performs a Fourier transform of the input, applies a learned linear transformation to the lower Fourier modes, and then returns to spatial space via inverse transform. We use the tensorized variant implemented in the `neuraloperator` package [43, 44], which applies Tucker factorization to the Fourier weights to reduce the number of parameters [45].

The FNO is trained using complex-valued transverse magnetization fields generated by the dMRI simulations as described in Section 2.2.3. Since the FNO implementation operates on regular Cartesian grids, the magnetization fields on the 160 µm × 160 µm area simulated by the SpinDoctor solver, which are originally defined on unstructured triangular meshes, are interpolated onto uniform grids prior to training. The real and imaginary components of the magnetization are interpolated onto square Cartesian grids containing 220 × 220 grid points (corresponding to a spacing of approximately 0.73 µm over the 160 µm × 160 µm domain). This grid size was empirically selected to balance spatial fidelity and computational efficiency, and was found to have negligible impact on signal accuracy (see Appendix B). This regular-grid representation ultimately enables meshfree inference directly from cell segmentation masks.

The FNO input comprises four channels:

1. The *x*- and *y*-coordinates of the grid points,
2. A diffusivity map, assigning each grid point the intracellular diffusivity, extracellular diffusivity, or zero at membrane locations,
3. A permeability map, where grid points corresponding to membrane boundaries are assigned the membrane permeability and all other locations are set to zero.

Compartment assignments (intracellular space, extracellular space, or membrane) are derived from the computational mesh used for the SpinDoctor simulations by assigning each grid point the label of the triangle in which it resides. During inference, however, compartment assignments are obtained directly from the cell segmentation masks, eliminating the need for mesh generation. The impact of this mesh-free inference strategy is evaluated in Appendix E.

The FNO outputs the complex-valued transverse magnetization field at the end of the diffusion-encoding sequence, represented by two real-valued channels corresponding to the real and imaginary components and defined on the same regular grid as the input. The dMRI signal is subsequently obtained by spatial integration of the predicted magnetization over the central 32 µm × 32 µm region, which represents the effective subdomain output used by the tiling framework. In preliminary experiments we found that predicting the full magnetization field yielded substantially higher accuracy than directly regressing the dMRI signal.

An example of a histology-derived label image, the corresponding diffusivity and permeability maps, and the reference magnetization field is shown in Fig. 4. Prior to training, each input and output channel is independently normalized by subtracting its mean and dividing by its standard deviation, where the normalization statistics are computed over the training set.

**Figure 4.**
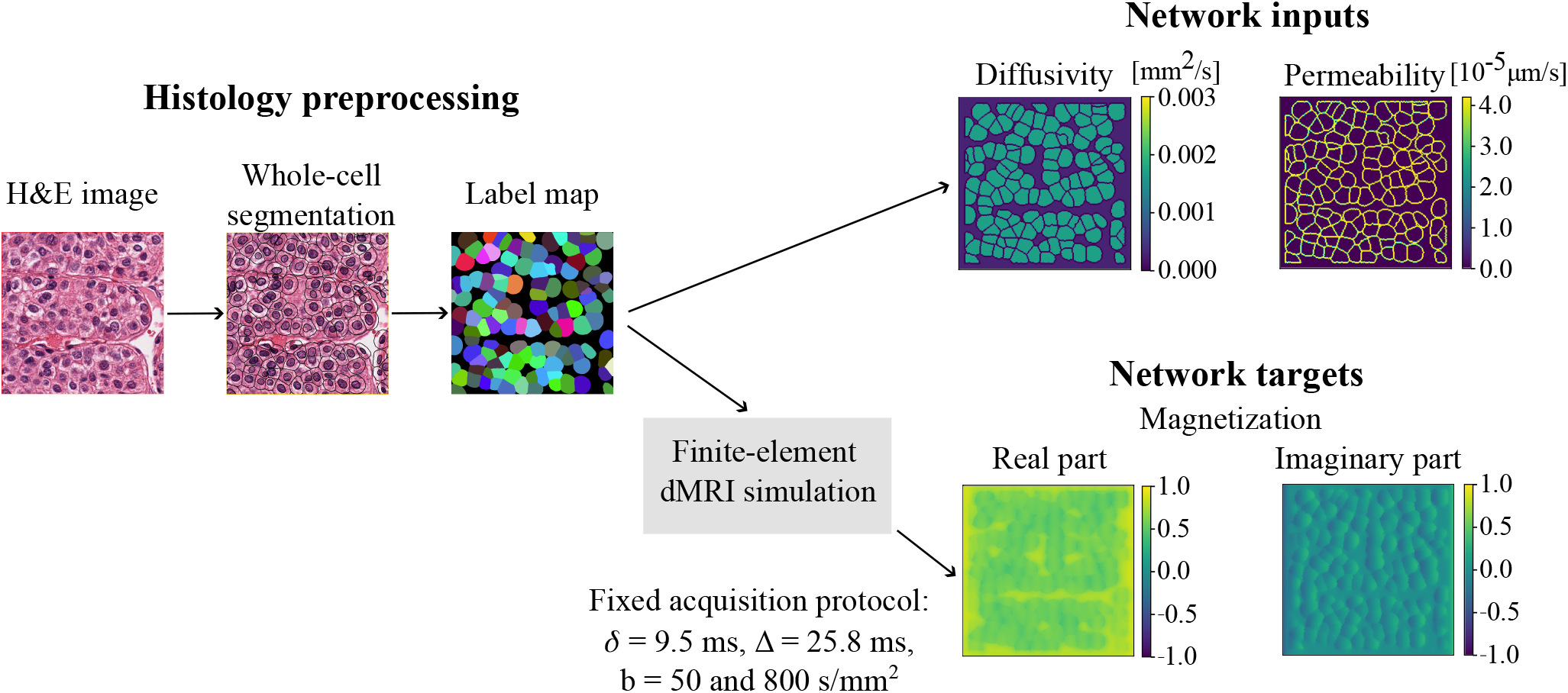
Example of the data-generation pipeline for a histology-derived patch. The H&E image is processed to obtain a whole-cell segmentation. The label map is converted into diffusivity and permeability maps, which form the FNO input together with spatial coordinates (not shown). The same label map is used to generate a finite-element mesh for dMRI simulation (SpinDoctor), producing the complex-valued transverse magnetization field (real and imaginary components) used as reference targets. Simulations were performed using a fixed PGSE diffusion-encoding protocol with gradient pulse duration *δ* = 9.5 ms, pulse separation Δ = 25.8 ms, and *b*-values of 50 and 800 s/mm^2^.

### 2.4. Training setup

To evaluate whether a neural operator can serve as a reusable model across different diffusion encoding strengths and biophysical assumptions, separate FNO models were trained for each diffusion weighting (*b* = 50 s/mm^2^ and *b* = 800 s/mm^2^) and dataset configuration (fixed-parameter and variable-parameter; see Section 2.2.3).

All models were trained to minimize the mean squared error between the predicted and reference simulated complex-valued magnetization fields.

Hyperparameters were selected via a grid search conducted on the validation set of the fixed-parameter dataset at *b* = 800 s/mm^2^. The following parameters were explored:

1. Number of truncated Fourier modes per spatial dimension: {32, 64, 128}
2. Number of hidden channels: {16, 32, 64}
3. Number of FNO layers: {3, 4, 5}
4. Learning rate: {0.01, 0.005, 0.001}
5. Step size of the StepLR scheduler: {5, 10}

All other training parameters were held constant: 50 training epochs, batch size of 32, Adam optimizer, StepLR learningrate scheduler with decay factor *γ* = 0.5, and weight decay of 10^−4^.

The configuration yielding the lowest validation loss consisted of 128 Fourier modes, 64 hidden channels, 5 layers, a learning rate of 0.01, and a StepLR step size of 5. This configuration was used for all datasets and diffusion weightings.

Model accuracy was evaluated using the signal magnitude obtained by spatially integrating the predicted complex transverse magnetization over the retained central 32 µm × 32 µm region, corresponding to the effective output region of each extended subdomain in the tiling framework. This evaluation therefore directly measures the accuracy relevant for inference across whole-slide images.

Training was conducted on a NVIDIA A100 GPU (40 GB memory). A single training run took 50 minutes.

### 2.5. Evaluation on diverse cell morphologies

The training dataset used in this work is derived from automatically segmented histology images, in which cellular geometries are predominantly round or elliptical as a consequence of the segmentation pipeline (nucleus detection followed by morphological expansion). To assess the ability of the trained FNO to generalize beyond these simplified morphologies, we evaluate its performance on external datasets containing manually annotated cells with substantially greater shape variability.

We consider two publicly available whole-cell segmentation datasets:

1. CytoNuke [46]: 83 manually annotated label images from H&E-stained head and neck squamous cell carcinoma tissue.
2. TissueNet [47]: a subset of 3904 manually annotated label images spanning multiple tissue types, acquired using fluorescence and mass-spectrometry imaging modalities.

Representative examples from both datasets are shown in Fig. 5. In contrast to the relatively regular cell shapes present in the training data (see Fig. 2), these examples exhibit pronounced morphological irregularity and a wide range of cell sizes, providing a challenging test of model generalization. Additionally, Fig. 6 displays the distribution of cell count and ratio of cell to extracellular space (c.f. Section 2.2.2). For the TissueNet dataset, there are samples with higher cell count and greater overall cell area compared to the training set, whereas the features of the CytoNuke data fall within the training distribution.

**Figure 5.**
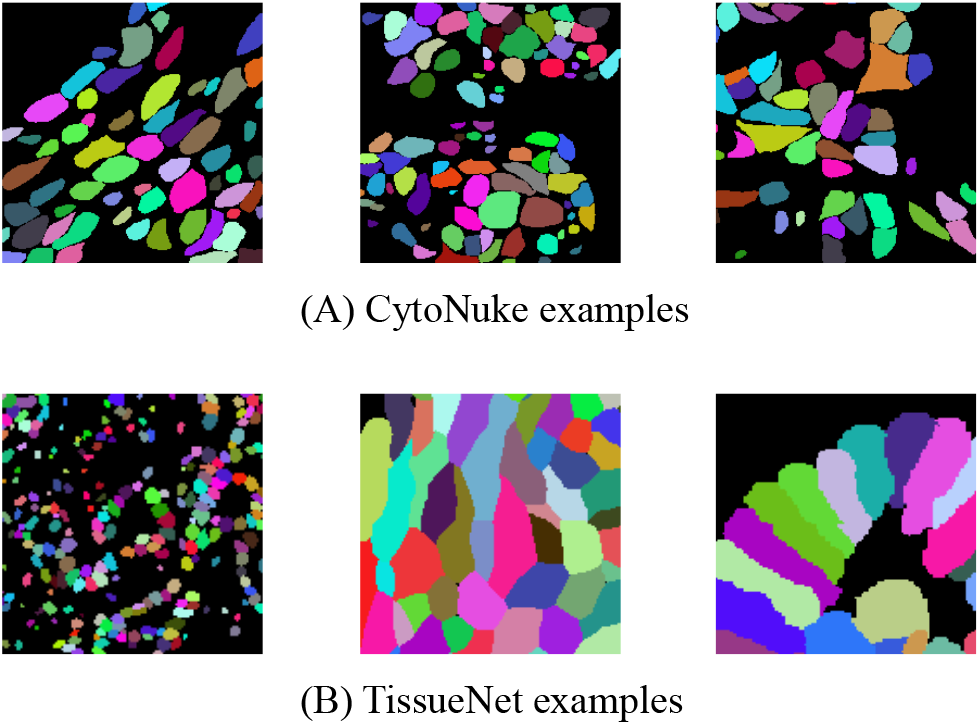
Representative cell segmentation masks from external datasets used to evaluate generalization. (A) CytoNuke: manually annotated cells from H&E-stained head and neck squamous cell carcinoma tissue. (B) TissueNet: manually annotated cells from fluorescence and mass-spectrometry data across diverse tissue types. Both datasets exhibit substantially more irregular and heterogeneous cell morphologies than those present in the automatically segmented training data.

**Figure 6.**
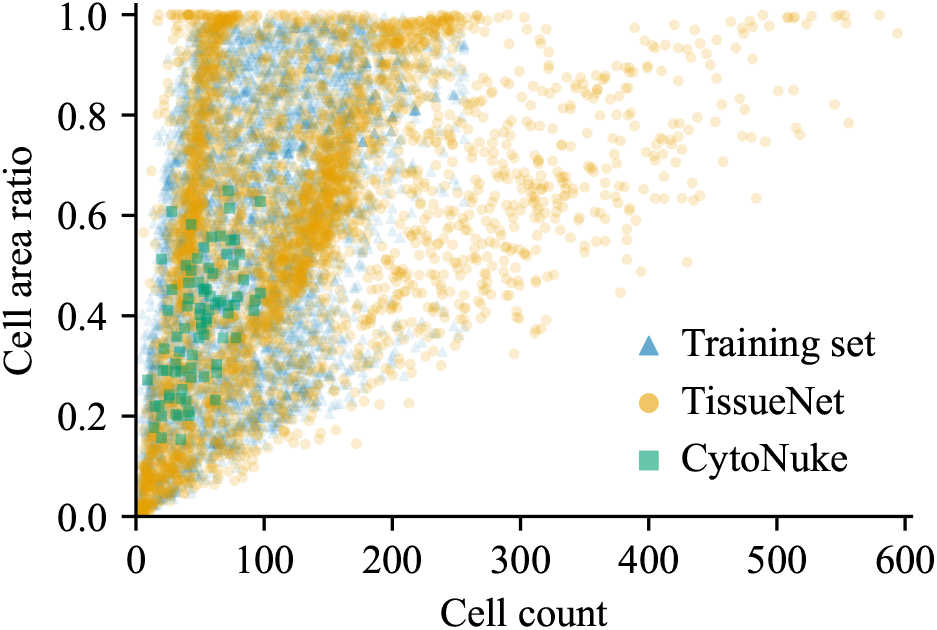
Distribution of cell count and cell area ratio across the training set and the two evaluation sets derived from the CytoNuke and TissueNet datasets. While the feature distribution of the Cytonuke data falls within that of the training set, the TissueNet data exhibits a much wider variety, in particular there are more samples with high cell count and high ratio of cells to extracellular space.

For both datasets, dMRI simulations are performed using same acquisition parameters and biophysical properties in the fixed-parameter training set, for two diffusion weightings: *b* = 50 and 800 s/mm^2^. The intracellular diffusivity is set to 0.001 mm^2^/s, the extracellular diffusivity to 0.003 mm ^2^/s, and the membrane permeability to 0 µm/s. Dataset-specific preprocessing details are provided in Appendix C.

Two FNO models, trained separately on the fixed-parameter datasets at *b* = 50 and *b* = 800 s/mm^2^ , are applied to the corresponding simulated datasets without additional finetuning. Model predictions are evaluated by computing the dMRI signal magnitude from the complex magnetization field within a central 32 µm × 32 µm region, consistent with the evaluation protocol used for the original test set.

### 2.6. Whole-slide inference and ADC map computation

The primary motivation of the proposed framework is to enable dMRI simulation directly from whole-slide histology images at spatial scales that are computationally inaccessible using conventional finite-element approaches. We therefore evaluate the framework by generating ADC maps across an entire histological whole-slide image.

A representative liver whole-slide image from the TCGA dataset (case ID: TCGA-UB-AA0U) was selected for this study. Following QIBA recommendations for liver dMRI [1], we targeted an in-plane spatial resolution of 2 mm × 2 mm per pixel. Since the present experiments are restricted to two-dimensional histology-derived microstructure, all inference and ADC estimation are performed in-plane.

Each ADC map pixel is represented by a grid of 63 × 63 extended subdomains, each contributing a central region of 32 µm × 32 µm, resulting in an effective in-plane pixel size of 2016 µm × 2016 µm. A rectangular region comprising 14 × 9 pixels was analyzed, corresponding to a physical extent of 28.224 mm × 18.144 mm and yielding a total of 126 pixels.

For each pixel, the complex transverse magnetization fields were predicted using two trained FNO models corresponding to diffusion weightings of *b*=50 s/mm^2^ and *b*=800 s/mm^2^ .The magnetization for the gradient direction along the *y*-axis was obtained by rotating both the input label images and the predicted magnetization fields by 90°. This operation is valid under the assumption of isotropic diffusion in both intra- and extracellular compartments. For each voxel, dMRI signals were computed by averaging the predicted magnetization over all subdomains contributing to the voxel, followed by averaging across gradient directions. ADC values were then estimated using a two-point logarithmic fit between the two diffusion weightings,

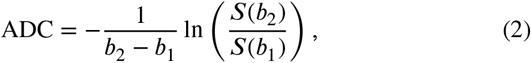

where *S*(*b*_1_) and*S*(*b*_2_) denote the direction-averaged signals at *b*_1_ = 50 s/mm^2^ and *b*_2_ = 800 s/mm^2^, respectively.

Because the proposed FNO reduces simulation times from months to hours, it becomes feasible to systematically investigate how segmentation assumptions and biophysical parameters influence image-scale dMRI measurements. We therefore generate ADC maps under two experimental conditions:

1. Segmentation parameter variation: Physical parameters were held fixed (extracellular diffusivity 0.003 mm ^2^/ s, intracellular diffusivity 0.001 mm ^2^/s, zero membrane permeability), while the cell expansion radius used during the segmentation with QuPath was varied between 1 and 7 µm. The expansion radius determines how far detected nuclei are radially enlarged to approximate the cytoplasm. Increasing the expansion radius therefore produces larger effective cell regions and reduces extracellular space. This experiment assesses the sensitivity of ADC estimates to segmentation choices.
2. Physical parameter sweep: Segmentation parameters were fixed (cell expansion radius 5 µm, detection threshold 0.4), while extracellular diffusivity ({0 001, 0 002, 0 003} mm /s), intracellular diffusivity ({0.0002, 0.001, 0.002, 0.003} mm /s), and membrane permeability ({0, 26, 50} µm/s) were varied, yielding 36 parameter combinations. This analysis illustrates how assumed biophysical properties influence macroscopic ADC values.

## 3. Results

### 3.1. Fourier Neural Operator prediction accuracy

We first evaluate whether the proposed neural operator can faithfully replace local finite-element simulations, since accurate prediction of subdomain magnetization is a prerequisite for whole-slide inference.

We evaluate the trained FNO models described in Section 2.4 on the test sets comprising 1000 samples each, for both fixed-parameter and variable-parameter datasets and for diffusion weightings of *b*=50 and 800 s/mm^2^ (see Section 2.2.3). Prediction accuracy is quantified using the relative error between the dMRI signal magnitudes obtained by spatially integrating the predicted and FEM-reference complex transverse magnetization fields over the central 32 µm × 32 µm region of each subdomain.

Quantitative results are reported in Table 1. At *b*=800 s/mm^2^, the FNO achieves a mean relative error of 0.43% on the fixed-parameter dataset and 0.34% on the variable-parameter dataset. At the lower diffusion weighting of *b*=50 s/mm^2^ , errors are substantially reduced, with mean relative errors of 0.03% for both datasets.

**Table 1.** Relative error statistics of the signal magnitude obtained by spatial integration of the predicted magnetization over the central 32 µm × 32 µm region for the four test sets. Each test set contains 1000 samples.

| Dataset | $b$-value<br>(s/mm<sup>2</sup>) | Mean<br>(%) | Median<br>(%) | Std.<br>(%) | Max<br>(%) |
| --- | --- | --- | --- | --- | --- |
| Fixed | 800 | 0.43 | 0.32 | 0.42 | 3.49 |
| Fixed | 50 | 0.03 | 0.02 | 0.03 | 0.20 |
| Variable | 800 | 0.34 | 0.23 | 0.35 | 2.42 |
| Variable | 50 | 0.03 | 0.02 | 0.03 | 0.19 |

The full distributions of relative errors are shown in Figure 13 in Appendix D. Across all datasets, the distributions are concentrated at low error values with only a small number of higher-error outliers.

Across all evaluated settings, the error distributions are narrow, with maximum relative errors remaining below 5%. At lower diffusion weighting, both mean and worst-case errors decrease by approximately an order of magnitude, indicating that the FNO accurately captures the magnetization under low diffusion encoding.

### 3.2 Generalization to diverse cell morphologies

Since whole-slide histology inevitably contains tissue configurations that differ from those observed during training, robust out-of-distribution generalization is essential for practical deployment of the proposed framework. We evaluated the FNO trained on the fixed-parameter dataset at *b*=800 s/mm2 on two external datasets, CytoNuke and TissueNet, which exhibit substantially more complex and heterogeneous cell morphologies than those observed during training (Section 2.5). No fine-tuning or retraining was performed.

Quantitative results for the relative error between the signal magnitudes obtained from the predicted and reference magnetization fields over the central 32 µm × 32 µm region are summarized in Table 2. Although a performance degradation is observed for the external datasets, the increase in error remains modest given the pronounced differences in cellular geometry, compartmental complexity, and spatial organization.

**Table 2.** Relative error statistics of the signal magnitude obtained by spatial integration of the predicted magnetization over the central 32 µm × 32 µm region for the external datasets. The FNO used here was trained on the fixed-parameter datasets at *b* = 800 and 50 s/mm^2^.

| Dataset | $b$-value<br>(s/mm<sup>2</sup>) | Mean<br>(%) | Median<br>(%) | Std.<br>(%) | Max<br>(%) |
| --- | --- | --- | --- | --- | --- |
| CytoNuke | 800 | 1.79 | 1.15 | 1.74 | 7.79 |
| CytoNuke | 50 | 0.03 | 0.02 | 0.03 | 0.20 |
| TissueNet | 800 | 1.37 | 0.81 | 1.51 | 12.88 |
| TissueNet | 50 | 0.08 | 0.05 | 0.09 | 0.61 |

The full distributions of relative errors are shown in igure 13 in Appendix D. Across all datasets, the distributions are concentrated at low error values with only a small number of higher-error outliers.

Figure 7 illustrates a comparison between the predicted reference magnetization fields from the TissueNet dataset. This example corresponds to a relatively challenging case within the dataset, with a relative signal error of 8.44% within the central 32 µm × 32 µm evaluation region (consistent with the upper range of errors reported in Table 2). Even in this higher-error case, the largest errors are localized to cell boundaries and very small compartments. Notably, the reference magnetization exhibits visibly jagged cell boundaries, which arise from geometric artifacts introduced during the meshing process used for the finite element simulations. In contrast, the FNO-predicted magnetization displays smoother interfaces, reflecting the underlying regular grid representation. Similar error patterns are observed in the in-distribution test set, indicating that prediction inaccuracies are primarily associated with regions of high spatial gradients.

**Figure 7.**
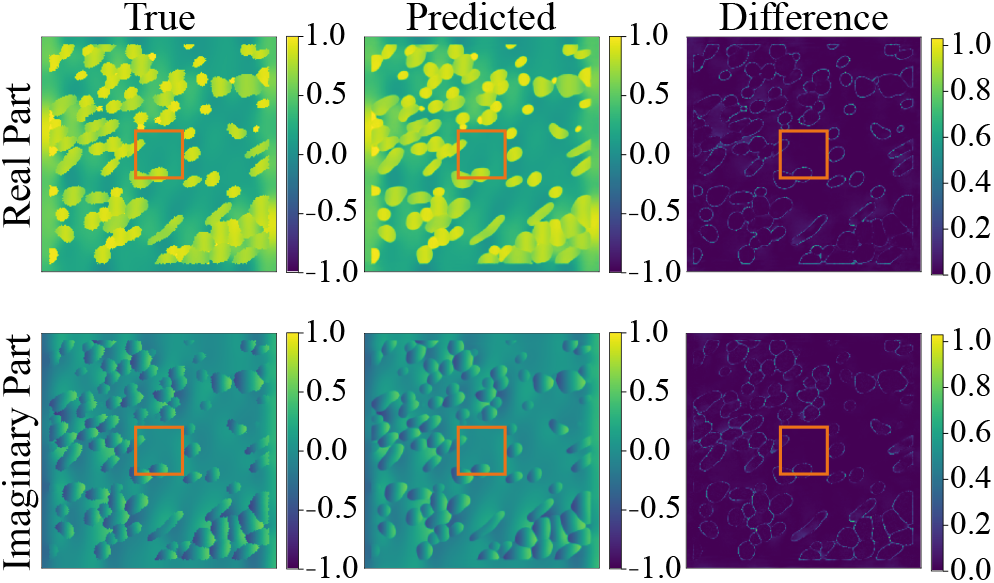
Reference magnetization simulated with SpinDoctor and corresponding FNO prediction for a sample from the TissueNet dataset at *b* = 800 s/mm^2^. The orange box outlines the central 32 µm × 32 µm region region used for signal evaluation. The relative error of the dMRI signal magnitude computed from this region is 8.44%. Discrepancies are most prominent near cell membranes, where the magnetization field exhibits sharp spatial discontinuities.

### 3.3. ADC maps based on whole-slide histology images

Having established local prediction accuracy and robustness across diverse morphologies, we next evaluate whether the proposed framework achieves its primary objective: practical dMRI simulation directly from whole-slide histology images.

By combining automatic cell segmentations with the trained FNO, dMRI metrics can be estimated directly from histological tissue architecture without performing numerical simulations for each subdomain. In the following experiments, we evaluate the computational requirements of this workflow and demonstrate its application to the generation of millimeter-scale ADC maps under varying cell segmentation and biophysical parameter settings.

Figure 8 shows the analyzed tissue region alongside a representative ADC map, illustrating how large-scale tissue structures are reflected in the resulting dMRI signal map.

**Figure 8.**
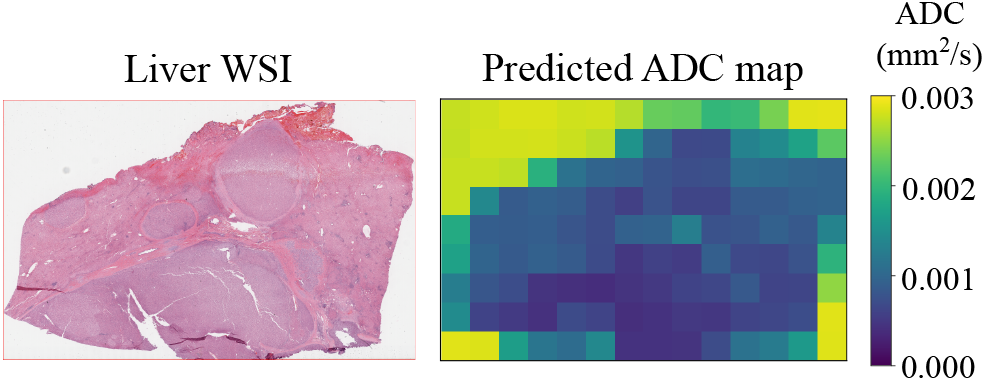
Generating ADC maps based on WSIs with the trained FNO. Left: Region of a whole-slide image used for analysis (case ID: TCGA-UB-AA0U) spanning 28,224 µm × 18,144 µm. Right: ADC map obtained with cell expansion radius of 5 µm and under fixed diffusivity and permeability parameters (extracellular diffusivity 0.003 mm^2^/s, intracellular diffusivity 0.001 mm^2^/s, zero permeability).

During inference, the trained FNO operates directly on parameter maps derived from cell segmentation masks and therefore does not require computational mesh generation or finite-element simulation. An evaluation of the impact of using label image–based inputs instead of mesh-derived inputs is provided in Appendix E, where only a modest increase in prediction error is observed.

#### 3.3.1. Computational efficiency

To assess whether the proposed neural operator enables practical dMRI simulation at clinically realistic scales, we evaluated the computational requirements of generating a complete ADC map for the whole-slide region shown in Figure 8, comprising 500,094 subdomains. In our previous work based on the subdomain framework [36], we performed finite-element simulations using SpinDoctor in two dimensions. On a dual-socket AMD EPYC 7543 system (64 cores, 128 threads, 1.1 TB RAM), a throughput of approximately 96 subdomains per hour was achieved using 64 concurrent processes, with further scaling limited by memory bandwidth. At this rate, generating a single ADC map based on a WSI for the present experiment would require approximately 217 days of computation for the finite element simulation alone, not accounting for the time required for mesh generation. Moreover, direct simulation of a full 2D domain of size 2016 µm × 2016 µm is not feasible in SpinDoctor due to memory constraints.

In contrast, using the trained FNO, preparation of label images of one ADC map for the input required 13 minutes on 30 CPU threads, while inference over all subdomains required 100 minutes on an NVIDIA A100 GPU (80 GB memory) with a batch size of 512. This >2600-fold acceleration transforms whole-slide histology-to-dMRI simulation from a computationally prohibitive task into a practical workflow.

#### 3.3.2. Effect of cell segmentation parameters

Because ADC maps based on WSIs can now be generated rapidly, it becomes feasible to systematically investigate how assumptions made during histological preprocessing influence image-scale dMRI measurements. We first investigated the sensitivity of ADC estimates to cell segmentation parameters. Specifically, the cell expansion radius was varied from 1 to 7 µm, while keeping the nucleus detection threshold fixed at 0.4 and holding all physical parameters constant (extracellular diffusivity *D*_ECS_ =0.003 mm ^2^/s, intracellular diffusivity *D*_ICS_=0.001 mm ^2^/s, and zero membrane permeability).

Figure 9 illustrates the effect of increasing the cell expansion radius on the resulting segmentation masks. Larger expansion radii produce larger effective cell regions and correspondingly smaller extracellular spaces.

**Figure 9.**
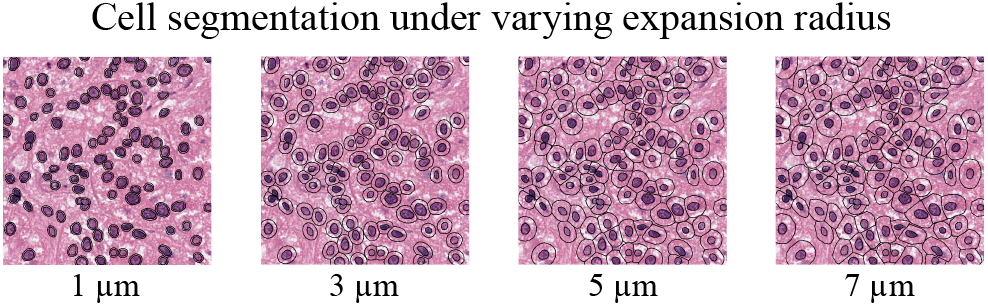
Illustration of the cell expansion radius used during the segmentation with QuPath. Beginning from the original H&E image patch (left), detected nuclei are radially expanded to approximate the whole cell. Segmentations generated with expansion radii of 1, 3, 5, and 7 µm are shown.

Figure 10 shows the resulting ADC maps for increasing expansion radius. A systematic decrease in ADC values is observed as the expansion radius increases, consistent with increased diffusion restriction arising from larger effective cell sizes. These results illustrate that the proposed framework enables quantitative assessment of uncertainty arising from histological preprocessing choices, an analysis that would be difficult to perform using conventional finite-element simulations at the scale of whole-slide images.

**Figure 10.**
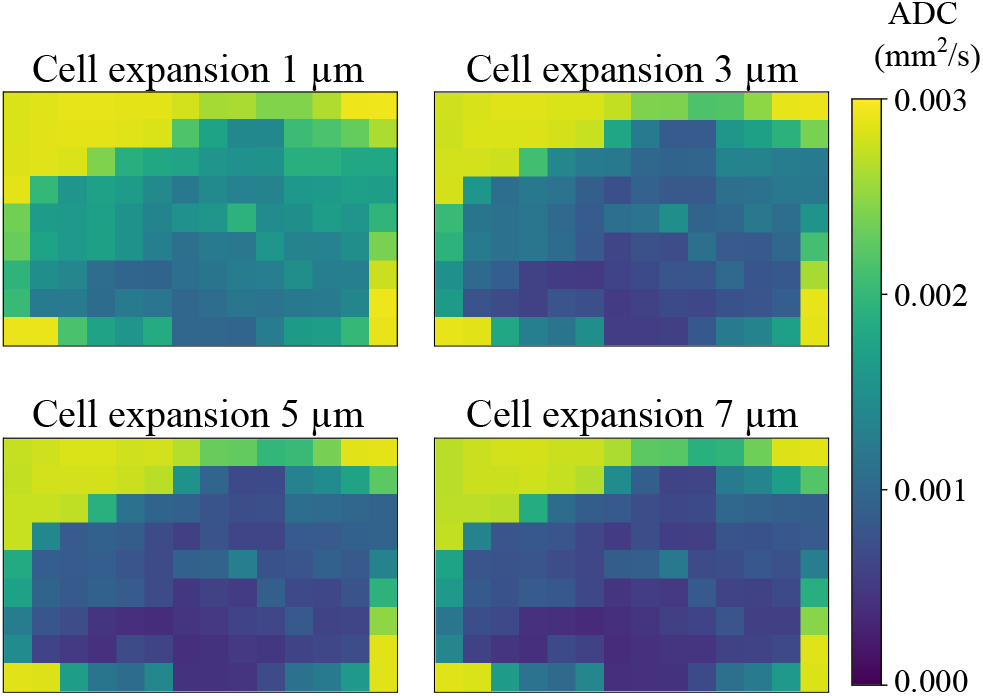
ADC maps computed from the same whole-slide image with varying cell expansion radius (1 to 7 µm) used during automatic cell segmentation. Extracellular diffusivity was fixed to 0.003 mm^2^/s, intracellular diffusivity to 0.001 mm^2^/s and permeability to zero. Increasing cell expansion systematically lowers ADC values due to more restricted diffusion in larger cells.

#### 3.3.3. Effect of diffusivity and permeability parameters

Next, we demonstrate that the framework can rapidly explore large biophysical parameter spaces by generating ADC maps from whole-slide histology images across 36 combinations of extracellular diffusivity, intracellular diffusivity, and membrane permeability. Figure 11 shows a representative subset of the results obtained with a fixed extracellular diffusivity of 0.003 mm^2^/s, intracellular diffusivities of 0.0002, 0.001, and 0.002 mm ^2^/s, and membrane permeabilities of 0, 26, and 50 µm/s. The complete set of ADC maps for all 36 parameter combinations is provided in Figure 14 in Appendix F.

**Figure 11.**
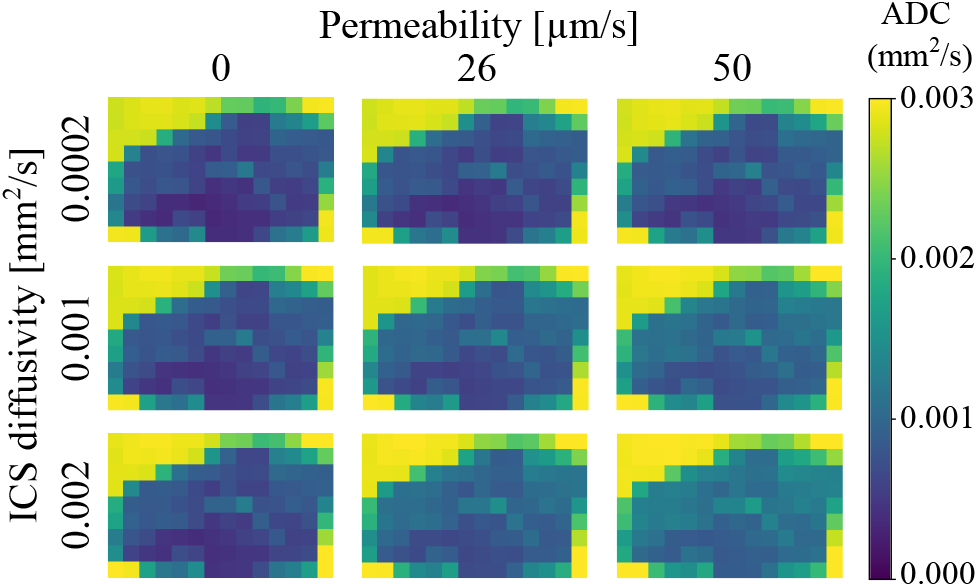
Representative ADC maps from the parameter sweep experiment. The extracellular diffusivity is fixed at 0.003 mm^2^/s, while intracellular diffusivity (rows) and membrane permeability (columns) are varied. Lower diffusivities lead to systematically lower ADC values, while higher permeability leads to increased ADC values. The complete set of results for all 36 parameter combinations is provided in Appendix F.

Decreasing either intracellular or extracellular diffusivity leads to uniformly lower ADC values across the tissue, reflecting reduced microscopic mobility. In contrast, increasing membrane permeability results in higher ADC values, consistent with enhanced exchange between compartments. Taken together, these experiments demonstrate that the proposed framework enables systematic investigation of how microscopic biophysical assumptions propagate to macroscopic dMRI measurements across entire histological sections.

## 4. Discussion

The present work demonstrates that large-scale histology-based dMRI simulation can be made practical by exploiting the locality of dMRI signal formation. Because spins explore only a limited neighborhood during a typical dMRI experiment, whole-slide simulations can be decomposed into a large number of local diffusion problems. By learning the mapping from local tissue architecture and biophysical parameters to the resulting transverse magnetization field, the proposed framework amortizes these repeated simulations and replaces computationally expensive finite-element solves with efficient neural network inference.

Prediction errors were predominantly localized to cell boundaries, where the magnetization field exhibits sharp spatial discontinuities. Accurately representing such transitions requires high spatial frequencies, which are intrinsically difficult to capture using the smooth approximations learned by neural operators. This challenge is further compounded by mesh-generation artifacts in the reference simulations, which introduce jagged and irregular cell boundaries. Despite these localized discrepancies, mean relative signal errors remained below 0.5% for the in-distribution datasets and below 2% for substantially different external morphologies, indicating only a minor impact on the aggregated dMRI signal.

The model also generalized well to datasets containing considerably more heterogeneous and irregular cell morphologies than those encountered during training, suggesting that the learned operator captures general relationships between tissue microstructure and dMRI signal formation rather than being strongly tied to specific cell shape distributions. From a practical perspective, cellular organization is often relatively consistent within a given tissue type. Consequently, extending the framework to new tissues would likely require only the generation of targeted simulation data representative of the relevant morphology distribution, followed by retraining or fine-tuning of the neural operator.

The achieved error levels should also be interpreted in the context of measurement uncertainty in clinical dMRI. Reported signal-to-noise ratios in abdominal dMRI correspond to relative noise levels of several percent [48, 49, 50, 51], substantially exceeding the errors observed in the present study. Consequently, the approximation error introduced by the FNO is unlikely to represent a major source of uncertainty in practical applications.

The primary advantage of the proposed framework lies in its computational efficiency. Replacing finite-element simulations with a trained neural operator reduces computation times from months to hours, enabling routine generation of ADC maps from whole-slide histology images. This makes it feasible to systematically investigate how dMRI measurements depend on underlying microstructural and biophysical parameters across large tissue regions, an analysis that would be impractical using direct numerical simulation alone.

A second practical advantage of the framework is that inference operates directly on regular-grid representations derived from histological cell segmentations. Unlike conventional finite-element and many Monte Carlo approaches, no computational mesh generation is required during deployment. This substantially simplifies the simulation pipeline and is particularly important for inference across whole-slide images, where generating meshes for hundreds of thousands of subdomains would itself become computationally burdensome.

In addition, the FNO operates on a substantially coarser discretization than the finite-element solver used to generate the training data (0.73 µm versus approximately 0.3 µm). Despite this reduction in resolution, Appendix B shows that increasing the number of grid points beyond 220×220 yields negligible improvements in signal accuracy. This observation indicates that accurate prediction of the final dMRI signal may not require the spatial discretization density needed for numerical solution of the underlying PDE. This suggests that the dMRI signal is comparatively insensitive to fine-scale variations in the magnetization field, at least for the acquisition settings considered in this study.

Beyond enabling large-scale inference, the computational efficiency of the framework opens opportunities for further acceleration. In our hyperparameter experiments, a range of network architectures achieved similar prediction accuracy, suggesting that smaller models may provide favorable trade-offs between computational cost and performance. Furthermore, because neighboring regions in wholeslide histology images often exhibit similar tissue architecture, it may be possible to reduce computation by evaluating only a subset of representative subdomains and estimating neighboring values through interpolation or aggregation schemes. Investigating such strategies could further improve the scalability of large-scale dMRI simulations.

Several limitations remain. First, the present implementation is restricted to two-dimensional simulations. Extension to three-dimensional microstructures is an important next step toward fully realistic histology-to-dMRI modeling.

Second, the current implementation is restricted to a single PGSE acquisition with fixed sequence parameters and neglects transverse relaxation by assuming *T*_2_ = ∞. Consequently, the trained FNO approximates only the transverse magnetization for this specific acquisition setting. Extending the framework to additional diffusion weightings, gradient waveforms, echo times, or relaxation effects would require either training separate neural operators or incorporating these acquisition and tissue parameters as additional inputs to a single conditional model. Although inference would remain computationally efficient once trained, generating the corresponding reference simulations as training data would introduce additional computational cost. Furthermore, in tissues where compartment-specific *T*_2_ differences substantially contribute to diffusion contrast, incorporating relaxation into the neural operator will be important for achieving realistic predictions. Extending the framework to arbitrary gradient waveforms, acquisition parameters, and additional relaxation mechanisms would substantially broaden its applicability and move toward a general-purpose surrogate for dMRI simulation.

Overall, the proposed framework demonstrates that neural operators can transform histology-based dMRI simulation from a computationally prohibitive numerical problem into a practical inference problem. By exploiting the locality and recurrence of dMRI signal formation, the method enables rapid estimation of diffusion measurements directly from whole-slide histology images and facilitates investigations of microstructure–signal relationships at spatial scales that were previously inaccessible.

## CRediT authorship contribution statement

**Iris A. Kohler:** Conceptualization, Data curation, Formal analysis, Investigation, Methodology, Software, Validation, Visualization, Writing - original draft, Writing - review & editing. **Oliver Gödicke:** Writing - review & editing. **Tristan A. Kuder:** Funding acquisition, Writing - review & editing. **Mark E. Ladd:** Funding acquisition, Writing - review & editing. **Jürgen Hesser:** Conceptualization, Funding acquisition, Project administration, Supervision, Writing - review & editing.

## Code and Data availability

The code is available at https://github.com/irisakohler/histology-to-dmri-neural-operator. The simulation data generated in this study is available upon request.

## Acknowledgment

The results shown here are in whole or part based upon data generated by the TCGA Research Network: https://www.cancer.gov/tcga.

## A. Appendix Whole-slide images used for training data generation

The case IDs of the 10 whole slide images from the TCGA-LIHC dataset that were used for the generation of the training, validation and test sets are shown in Table 3, along with the top left coordinates of the region that was extracted from each image.

**Table 3.** Top-left coordinates (in µm) of the 3150 µm × 3150 µm crop regions extracted from whole slide images used for data generation of training, validation and test set.

| Case ID | Top-left coordinates ($\mu\text{m}$) of crop region |
| --- | --- |
| TCGA-BC-A69I | 4000, 11,000 |
| TCGA-G3-A7M6 | 16,000, 12,000 |
| TCGA-YA-A8S7 | 12,500, 9500 |
| TCGA-CC-A1HT | 5000, 3100 |
| TCGA-CC-A3MA | 5500, 8500 |
| TCGA-K7-AAU7 | 7400, 5300 |
| TCGA-G3-A5SK | 14,200, 4500 |
| TCGA-DD-AAED | 2700, 5500 |
| TCGA-CC-A8HT | 6200, 6700 |
| TCGA-CC-5261 | 5700, 6200 |

## B. Appendix Grid spacing and interpolation strategy

The implementation of the FNO [43] requires input data to be defined on a regular Cartesian grid. However, SpinDoctor outputs the complex-valued magnetization field on a triangular mesh, where each mesh node stores a complex magnetization value. At compartment boundaries, nodes are duplicated, with each copy assigned a value corresponding to a different compartment, since the magnetization is discontinuous across membranes.

To prepare data for FNO training, we interpolate the magnetization from the mesh onto a regular 2D grid. We employ a nearest-neighbor interpolation strategy that respects compartment boundaries. Each grid point is first assigned a compartment label by identifying the triangle of the mesh in which it lies. The magnetization value at that grid point is then obtained from the nearest mesh node belonging to the same compartment. This corresponds to a Voronoi-type interpolation restricted to individual compartments and ensures that discontinuities across membranes are preserved.

The computational domain corresponds to a square region of size 160 µm × 160 µm. For a given grid size *N*, a uniform Cartesian grid with *N* × *N* points is constructed.

To evaluate the effect of the interpolation grid size, we compare the dMRI signal magnitude computed from the interpolated grid to the reference signal obtained directly from the mesh-based SpinDoctor simulation.

In SpinDoctor, the signal is computed by integrating the complex magnetization over the mesh using the finite element mass matrix. On the Cartesian grid, we approximate the spatial integral numerically using the trapezoidal rule.

We compute the relative signal magnitude error as

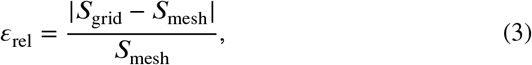

where *S*_grid_ is the dMRI signal computed from the grid and *S*_mesh_ the signal computed from the mesh. The evaluation is performed on the validation dataset consisting of 1000 samples.

We evaluate grid sizes ranging from 40 × 40 up to 800 × 800 on the four training datasets: the fixed-parameter and variable-parameter datasets at *b* = 50 and 800 s/mm^2^. Figure 12 shows the mean relative signal error for each dataset as a function of grid size.

**Figure 12.**
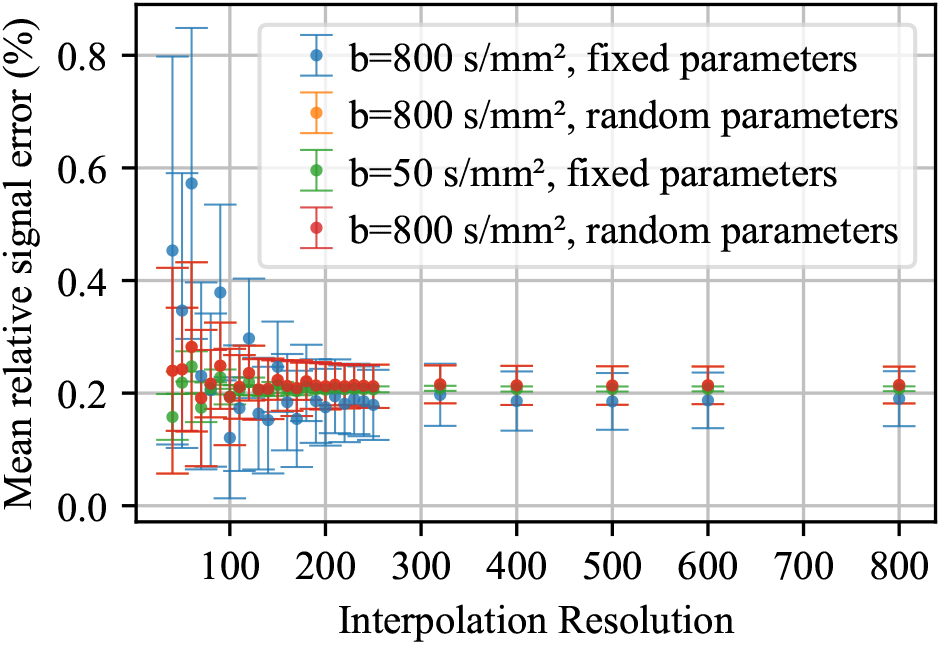
Mean relative signal error as a function of grid size for the fixed-parameter and variable-parameter datasets at *b* = 50 and 800 s/mm^2^.

**Figure 13.**
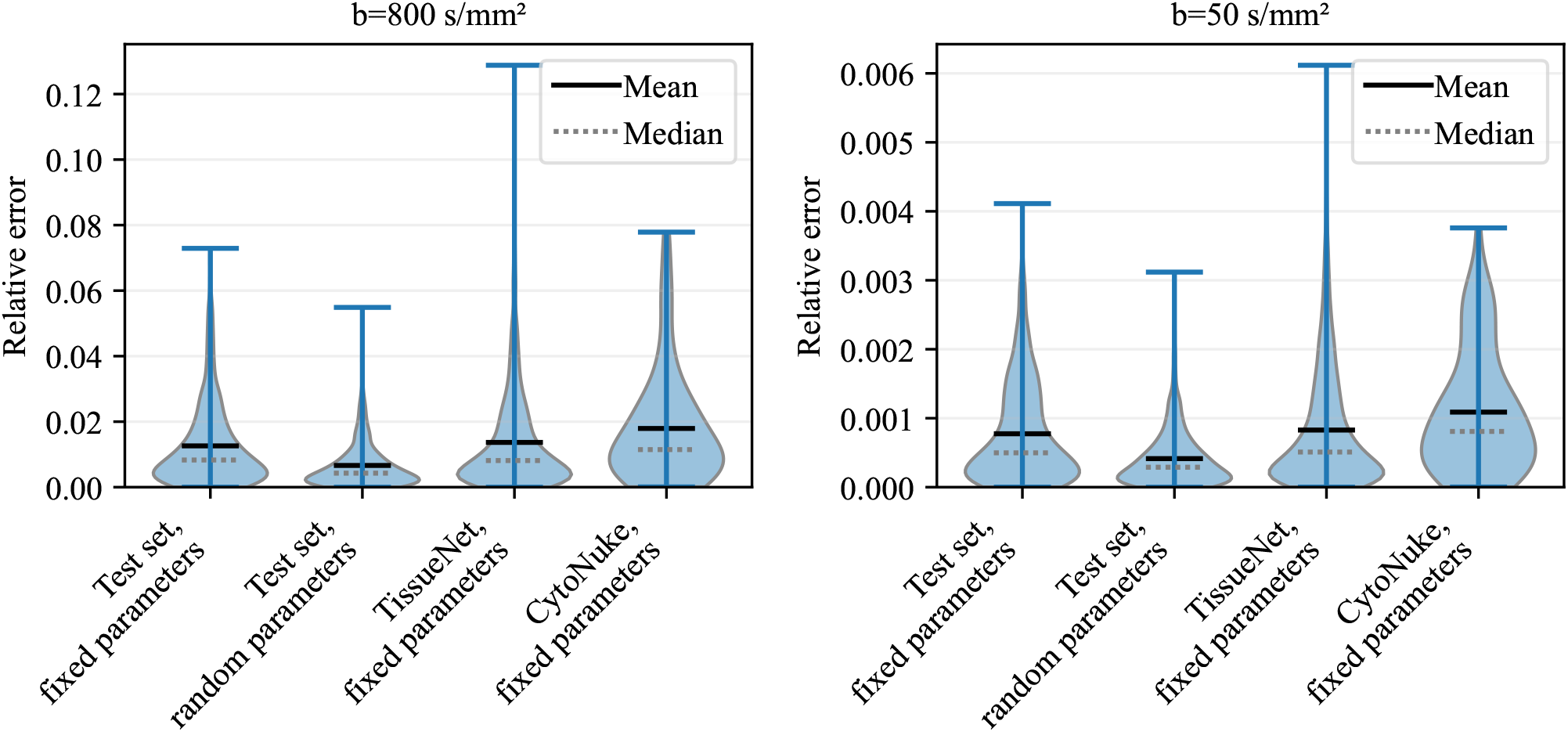
Distribution of relative signal errors across all evaluated datasets. The left panel shows results for *b* = 800 s/mm^2^ and the right panel for *b* = 50 s/mm^2^. In both cases, the distributions are strongly concentrated at low error values with narrow tails extending toward higher errors. The overall distribution shapes are similar across diffusion weightings, although error magnitudes are substantially lower for *b* = 50 s/mm^2^.

**Figure 14.**
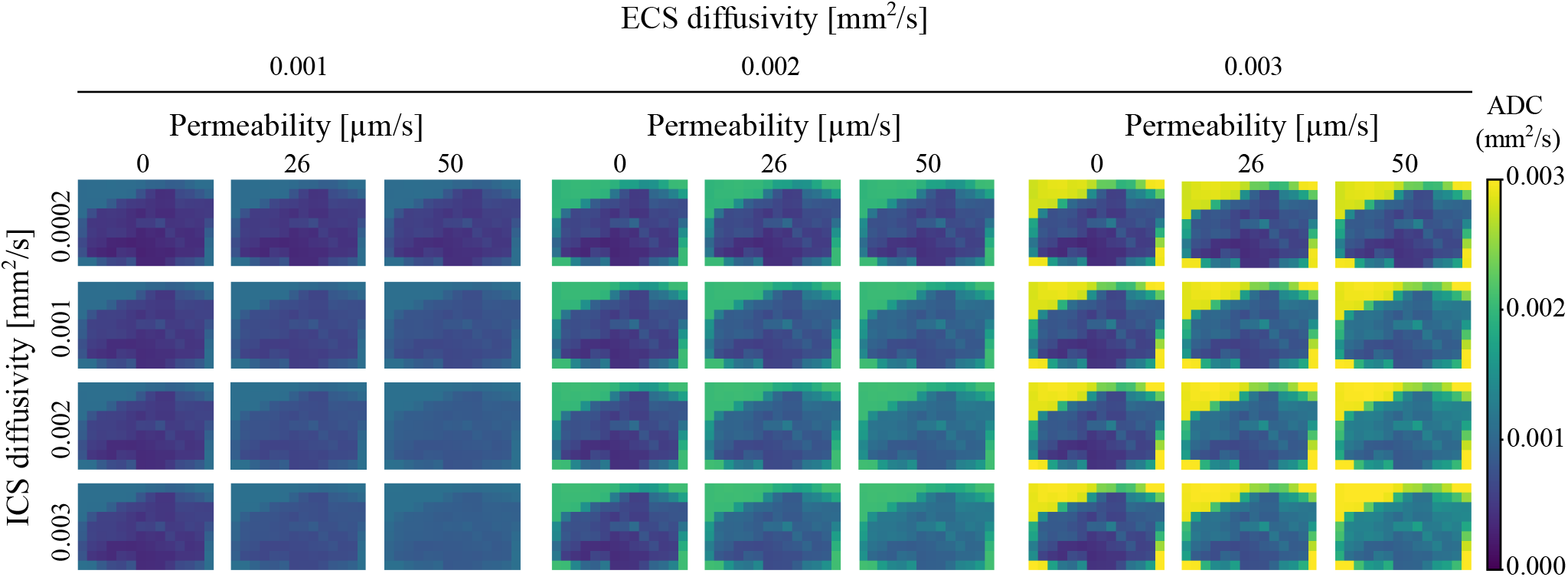
Complete parameter sweep for ADC maps derived from whole-slide histology images. ADC maps are shown for all combinations of extracellular (ECS) diffusivity ({0.001, 0.002, 0.003} mm^2^/s), intracellular (ICS) diffusivity ({0.0002, 0.001, 0.002, 0.003} mm^2^/s), and membrane permeability ({0, 26, 50} µm/s). Lower diffusivities generally result in reduced ADC values, whereas higher membrane permeabilities increase ADC by facilitating exchange between intracellular and extracellular compartments.

For coarse grids, the relative signal error is dominated by is discretization effects and can exceed 0.5%. As the grid refined, the error rapidly decreases and stabilizes around 0.2% for grid sizes above approximately 100 × 100.

Notably, further increasing the grid size beyond 200 × 200 does not lead to a reduction in error. We attribute this behavior to a geometric mismatch between the mesh and the regular grid. The triangular mesh does not cover a perfectly rectangular domain, whereas the Cartesian grid uniformly samples the full bounding box. As a result, the grid-based integration includes regions near the boundary that are not part of the original mesh domain.

To verify this hypothesis, we repeated the integration while restricting it to grid points located strictly inside the mesh. In this case, the relative signal error continued to decrease as the grid was refined, reaching values as low as 0.015% for a grid size of 1600 × 1600.

Based on the above analysis, we select a grid size of 220 × 220 for all experiments. At this grid size, the relative signal error is stable and close to its asymptotic value.

## C. Appendix Processing of TissueNet and CytoNuke

To evaluate the generalization capability of the trained FNO to diverse and previously unseen cell morphologies, we utilize two publicly available datasets with manually segmented whole-cell annotations: CytoNuke [46] and TissueNet [47]. These datasets feature a wide range of cellular shapes and spatial configurations that differ significantly from the predominantly round or elliptical cells in our training data, which were derived from automatically segmented histology images.

To ensure comparability with the training domain, all images are preprocessed to match the input field of view (FOV) of 150 µm × 150 µm. This is consistent with the simulation setup used during training, where each sample corresponds to a 150 µm patch.

### C.1. TissueNet

TissueNet provides over 4000 microscopy images with pixel-level whole-cell segmentations. Images span several modalities (fluorescence, mass spectrometry) and resolutions, with pixel sizes ranging from 0.365 to 1.0 µm, corresponding to FOVs between approximately 93 µm × 93 µm and 512 µm × 512 µm.

To adapt these images to the required FOV and resolution, we apply the following preprocessing steps:

1. Images with FOV < 150 µm × 150 µm are discarded.
2. From the remaining images, random crops of size 150 µm × 150 µm are extracted.
3. Each cropped image is then upscaled to a resolution of 1500 ×1500 pixels.
4. Images containing no segmented cells are discarded.

After filtering and preprocessing, 3844 images remain. Surface meshes are generated using the same pipeline as for the training, validation, and test data, with the `facet_size` parameter set to 3.0.

#### C.2. CytoNuke

CytoNuke consists of high-resolution single-cell segmentations across 83 images, each with original dimensions of 256 × 256 pixels and a reported pixel size of 0.5 µm, yielding a native FOV of 128 µm × 128 µm, slightly smaller than the FNO input domain.

To ensure consistency with the training domain, the following preprocessing is applied:

1. All images are upscaled to a resolution of 1500 × 1500 pixels.
2. During this process, a pixel spacing of 0.1 µm is assumed, resulting in an effective FOV of 150 µm × 150 µm, although this does not reflect the true physical scale of the dataset.

While this assumption introduces a discrepancy in absolute scale, it preserves the geometric complexity of the cell shapes, which is the primary factor of interest for evaluating out-of-distribution generalization.

Surface meshes are subsequently generated using the same pipeline as for the training data, with the `facet_size` parameter set to 3.0.

## D. Appendix Distribution of relative signal errors

Figure 13 shows the distribution of relative signal errors for all evaluated datasets at diffusion weightings of = 800 and 50 s/mm^2^ .The violin plots complement the summary statistics reported in Tables 1 and 2 by visualizing the full error distributions.

## E. Appendix Impact of mesh-free input representation

The cell boundaries represented in the computational mesh do not perfectly align with those in the original label images. This discrepancy arises because the meshing procedure introduces geometric approximations, leading to slightly jagged and irregular cell interfaces. In the results reported above, the FNO input parameter maps were derived directly from the mesh to ensure consistency with the numerical simulations.

However, mesh generation is computationally expensive and therefore impractical for applications requiring fast inference. To assess the impact of this limitation, we additionally evaluate the prediction error when the input parameter maps are instead derived directly from the label images, bypassing the meshing step. Quantitative results are summarized in Table 4.

**Table 4.**
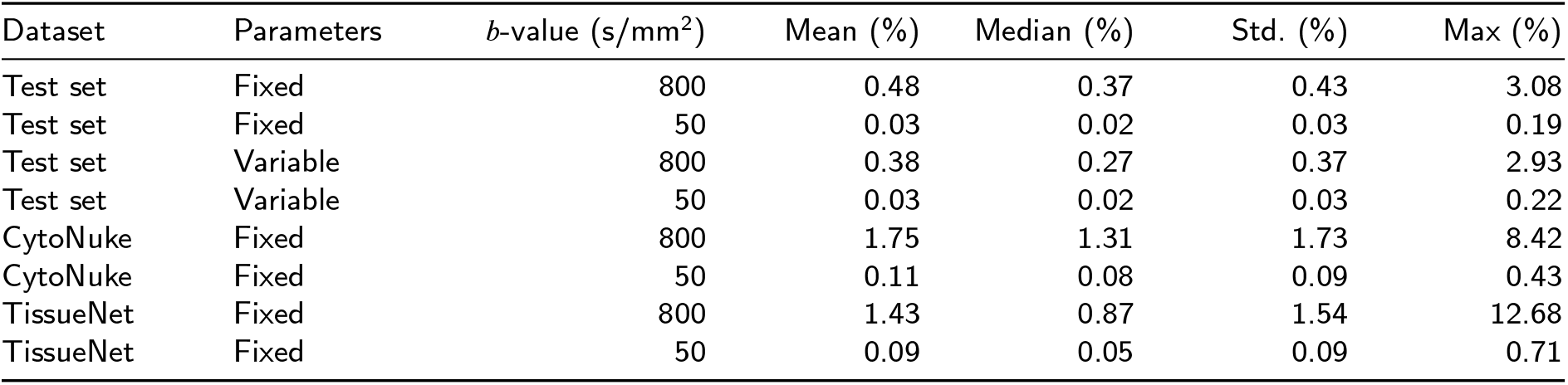
Relative error statistics of the signal in the central 32 µm × 32 µm region of the predicted magnetization when input parameter maps are derived from label images rather than the computational mesh.

Overall, the use of label image–based parameter maps leads to a modest increase in error, particularly at higher *b*-values and for datasets with more complex morphologies. Nevertheless, the errors remain low across all settings, indicating that the FNO is robust to moderate geometric inconsistencies between training and inference representations.

## F. Appendix Complete parameter sweep for ADC maps derived from whole-slide histology images

Figure 14 presents the complete set of ADC maps derived from WSIs generated for all 36 combinations of extracellular diffusivity, intracellular diffusivity, and membrane permeability described in Section 2.6. These results complement the representative subset shown in Figure 11 and provide a comprehensive overview of the influence of the investigated biophysical parameters on the resulting ADC maps.

## References

[1] A. Shukla-Dave, N. A. Obuchowski, T. L. Chenevert, S. Jambawalikar, L. H. Schwartz, D. Malyarenko, W. Huang, S. M. Noworolski, R. J. Young, M. S. Shiroishi, H. Kim, C. Coolens, H. Laue, C. Chung, M. Rosen, M. Boss, E. F. Jackson, Quantitative imaging biomarkers alliance (QIBA) recommendations for improved precision of DWI and DCE-MRI derived biomarkers in multicenter oncology trials, Journal of Magnetic Resonance Imaging 49 (2019) e101–e121.

[2] E. Panagiotaki, M. G. Hall, H. Zhang, B. Siow, M. F. Lythgoe, D. C. Alexander, High-fidelity meshes from tissue samples for diffusion MRI simulations, in: Medical Image Computing and Computer-Assisted Intervention – MICCAI 2010, Springer Berlin Heidelberg, Berlin, Heidelberg, 2010, pp. 404–411.

[3] H. H. Lee, K. Yaros, J. Veraart, J. L. Pathan, F. X. Liang, S. G. Kim, D. S. Novikov, E. Fieremans, Along-axon diameter variation and axonal orientation dispersion revealed with 3D electron microscopy: implications for quantifying brain white matter microstructure with histology and diffusion MRI, Brain Structure and Function 224 (2019) 1469–1488.

[4] A. Grigoriou, C. Macarro, M. Palombo, D. Navarro-Garcia, A. K. Voronova, K. Bernatowicz, I. Barba, A. Escriche, E. Greco, M. Abad, S. Simonetti, G. Serna, R. Mast, X. Merino, N. Roson, M. Escobar, M. Vieito, P. Nuciforo, R. Toledo, E. Garralda, R. Sala-Llonch, E. Fieremans, D. S. Novikov, R. Perez-Lopez, F. Grussu, Histology-informed microstructural diffusion simulations for MRI cancer char-acterisation—the Histo-µSim framework, Communications Biology 8 (2025). Art. no. 1695.

[5] R. Gardier, A. Savoy, J. L. V. Haro, G. Girard, E. J. Canales-Rodriguez, E. Fischi-Gomez, A. Hertanu, I. O. Jelescu, J. Rafael-Patino, J. P. Thiran, Comparing Steam and PGSE diffusion MRI signal of rat lymph nodes using in-silico simulations, in: 2024 IEEE International Symposium on Biomedical Imaging (ISBI), IEEE Computer Society, 2024, pp. 1–5. doi:10.1109/ISBI56570.2024.10635470.

[6] G. Buizza, C. Paganelli, F. Ballati, S. Sacco, L. Preda, A. Iannalfi, D. C. Alexander, G. Baroni, M. Palombo, Improving the characterization of meningioma microstructure in proton therapy from conventional apparent diffusion coefficient measurements using Monte Carlo simulations of diffusion MRI, Medical Physics 48 (2021) 1250–1261.

[7] J. Bates, I. Teh, D. McClymont, P. Kohl, J. E. Schneider, V. Grau, Monte Carlo simulations of diffusion weighted MRI in myocardium: Validation and sensitivity analysis, IEEE Transactions on Medical Imaging 36 (2017) 1316–1325.

[8] M. Lashgari, N. Ravikumar, I. Teh, J. R. Li, D. L. Buckley, J. E. Schneider, A. F. Frangi, Three-dimensional micro-structurally informed in silico myocardium—towards virtual imaging trials in cardiac diffusion weighted MRI, Medical Image Analysis 82 (2022). Art. no. 102592.

[9] J. L. Villarreal-Haro, R. Gardier, E. J. Canales-Rodríguez, E. Fischi-Gomez, G. Girard, J. P. Thiran, J. Rafael-Patiño, CACTUS: a computational framework for generating realistic white matter microstructure substrates, Frontiers in Neuroinformatics 17 (2023). Art. no. 1208073.

[10] K. Ginsburger, F. Matuschke, F. Poupon, J.-F. Mangin, M. Axer, C. Poupon, MEDUSA: A GPU-based tool to create realistic phantoms I.A. Kohler et al.: Preprint submitted to Elsevier Page 15 of 17 of the brain microstructure using tiny spheres, NeuroImage 193 (2019) 10–24.

[11] H. H. Lee, E. Fieremans, D. S. Novikov, Realistic Microstructure Simulator (RMS): Monte Carlo simulations of diffusion in three-dimensional cell segmentations of microscopy images, Journal of Neuroscience Methods 350 (2021). Art. no. 109018.

[12] J. Rafael-Patino, D. Romascano, A. Ramirez-Manzanares, E. J. Canales-Rodríguez, G. Girard, J. P. Thiran, Robust Monte-Carlo simulations in Diffusion-MRI: Effect of the substrate complexity and parameter choice on the reproducibility of results, Frontiers in Neuroinformatics 14 (2020). Art. no. 8.

[13] L. Kerkelä, F. Nery, M. Hall, C. Clark, Disimpy: A massively parallel Monte Carlo simulator for generating diffusion-weighted mri data in python, Journal of Open Source Software 5 (2020). Art. no. 2527.

[14] A. Aghaeifar, S. Mueller, K. Scheffler, SpinWalk: A Monte Carlo simulator for MR-signal formation in inhomogeneous tissue, Imaging Neuroscience 3 (2025). Art. no. imag_a_00533.

[15] J. R. Li, V. D. Nguyen, T. N. Tran, J. Valdman, C. B. Trang, K. V. Nguyen, D. T. S. Vu, H. A. Tran, H. T. A. Tran, T. M. P. Nguyen, SpinDoctor: A MATLAB toolbox for diffusion MRI simulation, NeuroImage 202 (2019). Art. no. 116120.

[16] J. Xu, S. P. Devan, D. Shi, A. Pamulaparthi, N. Yan, Z. Zu, D. S. Smith, K. D. Harkins, J. C. Gore, X. Jiang, MATI: A GPU-accelerated toolbox for microstructural diffusion MRI simulation and data fitting with a graphical user interface, Magnetic Resonance Imaging 122 (2025). Art. no. 110428.

[17] Y. Jing, I. E. Magnin, C. Frindel, Monte Carlo simulation of water diffusion through cardiac tissue models, Medical Engineering and Physics 120 (2023). Art. no. 104013.

[18] J. Rafael-Patino, G. Girard, R. Truffet, M. Pizzolato, E. Caruyer, J.-P. Thiran, The diffusion-simulated connectivity (DiSCo) dataset, Data in Brief 38 (2021). Art. no. 107429.

[19] C. H. Yeh, B. Schmitt, D. L. Bihan, J. R. Li-Schlittgen, C. P. Lin, C. Poupon, Diffusion Microscopist Simulator: A general Monte Carlo simulation system for diffusion magnetic resonance imaging, PLoS ONE 8 (2013). Art. no. e76626.

[20] J. N. Rose, S. Nielles-Vallespin, P. F. Ferreira, D. N. Firmin, A. D. Scott, D. J. Doorly, Novel insights into in-vivo diffusion tensor cardiovascular magnetic resonance using computational modeling and a histology-based virtual microstructure, Magnetic Resonance in Medicine 81 (2019) 2759–2773.

[21] L. Wang, Y. Hong, Y. B. Qin, X. Y. Cheng, F. Yang, J. Yang, Y. M. Zhu, Connecting macroscopic diffusion metrics of cardiac diffusion tensor imaging and microscopic myocardial structures based on simulation, Medical Image Analysis 77 (2022). Art. no. 102325.

[22] D. Karimi, S. K. Warfield, Diffusion MRI with machine learning, Imaging Neuroscience 2 (2024) 1–55.

[23] N. Naughton, S. Cahoon, B. Sutton, J. G. Georgiadis, Accelerated, physics-inspired inference of skeletal muscle microstructure from diffusion-weighted MRI, IEEE Transactions on Medical Imaging (2024).

[24] N. Kovachki, Z. Li, B. Liu, K. Azizzadenesheli, K. Bhattacharya, A. Stuart, A. Anandkumar, Neural operator: learning maps between function spaces with applications to pdes, Journal of Machine Learning Research 24 (2023).

[25] J. Berner, M. Liu-Schiaffini, J. Kossaifi, V. Duruisseaux, B. Bonev, K. Azizzadenesheli, A. Anandkumar, Principled approaches for extending neural architectures to function spaces for operator learning (2025).

[26] K. Azizzadenesheli, N. Kovachki, Z. Li, M. Liu-Schiaffini, J. Kossaifi, A. Anandkumar, Neural operators for accelerating scientific simulations and design (2023).

[27] D. N. Tanyu, J. Ning, T. Freudenberg, N. Heilenkötter, A. Rademacher, U. Iben, P. Maass, Deep learning methods for partial differential equations and related parameter identification problems, 2023. doi:10.1088/1361-6420/ace9d4H.

[28] U. Subedi, A. Tewari, Annual review of statistics and its application operator learning: A statistical perspective 63 (2026) 48.

[29] Z. Li, N. Kovachki, K. Azizzadenesheli, B. Liu, K. Bhattacharya, A. Stuart, A. Anandkumar, Fourier Neural Operator for parametric partial differential equations (2020).

[30] G. Wen, Z. Li, Q. Long, K. Azizzadenesheli, A. Anandkumar, S. M. Benson, Real-time high-resolution CO2 geological storage prediction using nested Fourier neural operators, Energy and Environmental Science 16 (2023) 1732–1741.

[31] T. Kurth, S. Subramanian, P. Harrington, J. Pathak, M. Mardani, D. Hall, A. Miele, K. Kashinath, A. Anandkumar, FourCastNet: Accelerating global high-resolution weather forecasting using adaptive Fourier Neural Operators, in: Proceedings of the Platform for Advanced Scientific Computing Conference, PASC 2023, Association for Computing Machinery, Inc, 2023. doi:10.1145/3592979.3593412.

[32] B. Li, H. Wang, S. Feng, X. Yang, Y. Lin, Solving seismic wave equations on variable velocity models with Fourier Neural Operator, IEEE Transactions on Geoscience and Remote Sensing 61 (2023).

[33] S. Guan, K. T. Hsu, P. V. Chitnis, Fourier neural operator network for fast photoacoustic wave simulations, Algorithms 16 (2023).

[34] W. Peng, S. Qin, S. Yang, J. Wang, X. Liu, L. L. Wang, Fourier Neural Operator for real-time simulation of 3D dynamic urban microclimate, Building and Environment 248 (2024).

[35] T. Zhou, X. Wan, D. Z. Huang, Z. Li, Z. Peng, A. Anandkumar, J. F. Brady, P. W. Sternberg, C. Daraio, AI-aided geometric design of anti-infection catheters, Technical Report, 2024. URL: https://www.science.org.

[36] I. A. Kohler, L. Zheng, T. A. Kuder, O. Gödicke, M. E. Ladd, J. Hesser, Large-domain histology-based diffusion mri simulation via independent local simulations, bioRxiv (2026).

[37] B. J. Erickson, S. Kirk, Y. Lee, O. Bathe, M. Kearns, C. Gerdes, K. Rieger-Christ, J. Lemmerman, The Cancer Genome Atlas Liver Hepatocellular Carcinoma Collection (TCGA-LIHC) (Version 5) [Data set], https://doi.org/10.7937/K9/TCIA.2016.IMMQW8UQ, 2016. doi:10.7937/K9/TCIA.2016.IMMQW8UQ, The Cancer Imaging Archive.

[38] P. Bankhead, M. B. Loughrey, J. A. Fernández, Y. Dombrowski, D. G. McArt, P. D. Dunne, S. McQuaid, R. T. Gray, L. J. Murray, H. G. Coleman, J. A. James, M. Salto-Tellez, P. W. Hamilton, QuPath: Open source software for digital pathology image analysis, Scientific Reports 7 (2017). Art. no. 16878.

[39] U. Schmidt, M. Weigert, C. Broaddus, G. Myers, Cell detection with star-convex polygons, in: Medical Image Computing and Computer Assisted Intervention – MICCAI 2018, volume 11071 LNCS, Springer Verlag, 2018, pp. 265–273. doi:10.1007/978-3-030-00934-2_30.

[40] E. Fokkinga, J. A. Hernandez-Tamames, A. Ianus, M. Nilsson, C. M. Tax, R. Perez-Lopez, F. Grussu, Advanced diffusion-weighted MRI for cancer microstructure assessment in body imaging, and its relationship with histology, Journal of Magnetic Resonance Imaging 60 (2023) 1278–1304.

[41] F. Grussu, K. Bernatowicz, I. Casanova-Salas, N. Castro, P. Nuciforo, J. Mateo, I. Barba, R. Perez-Lopez, Diffusion MRI signal cumulants and hepatocyte microstructure at fixed diffusion time: Insights from simulations, 9.4T imaging, and histology, Magnetic Resonance in Medicine 88 (2022) 365–379.

[42] R. Gardier, J. L. V. Haro, E. J. Canales-Rodríguez, I. O. Jelescu G. Girard, J. Rafael-Patiño, J. P. Thiran, Cellular Exchange Imaging (CEXI): Evaluation of a diffusion model including water exchange in cells using numerical phantoms of permeable spheres, Magnetic Resonance in Medicine 90 (2023) 1625–1640.

[43] J. Kossaifi, N. Kovachki, Z. Li, D. Pitt, M. Liu-Schiaffini, V. Du-ruisseaux, R. J. George, B. Bonev, K. Azizzadenesheli, J. Berner, A. Anandkumar, A library for learning neural operators, arXiv preprint arXiv:2412.10354 (2025).

[44] V. Duruisseaux, J. Kossaifi, A. Anandkumar, Fourier neural operators explained: A practical perspective (2026).

[45] A. Tran, A. Mathews, L. Xie, C. S. Ong, Factorized Fourier Neural Operators (2021).

[46] J. Raufeisen, K. Xie, F. Hörst, T. Braunschweig, J. Li, J. Kleesiek, R. Röhrig, J. Egger, B. Leibe, F. Hölzle, A. Hermans, B. Puladi, Cyto R-CNN and CytoNuke dataset: Towards reliable whole-cell segmentation in bright-field histological images (2024).

[47] N. F. Greenwald, G. Miller, E. Moen, A. Kong, A. Kagel, T. Dougherty, C. C. Fullaway, B. J. McIntosh, K. X. Leow, M. S. Schwartz, C. Pavelchek, S. Cui, I. Camplisson, O. Bar-Tal, J. Singh, M. Fong, G. Chaudhry, Z. Abraham, J. Moseley, S. Warshawsky, E. Soon, S. Greenbaum, T. Risom, T. Hollmann, S. C. Bendall, L. Keren, W. Graf, M. Angelo, D. V. Valen, Whole-cell segmentation of tissue images with human-level performance using large-scale data annotation and deep learning, Nature Biotechnology 40 (2022) 555–565.

[48] S. D. Vasylechko, A. Tsai, O. Afacan, S. Kurugol, Self-supervised denoising diffusion probabilistic models for abdominal dw-mri, Magnetic Resonance in Medicine 94 (2025) 1284–1300.

[49] Q. Chen, S. Fang, Y. Yuchen, R. Li, R. Deng, Y. Chen, D. Ma, H. Lin, F. Yan, Clinical feasibility of deep learning reconstruction in liver diffusion-weighted imaging: Improvement of image quality and impact on apparent diffusion coefficient value, European Journal of Radiology 168 (2023).

[50] D. Zhao, X. Kong, K. Yang, J. Wan, Z. Liu, F. Pan, P. Sun, C. Zheng, L. Yang, Deep learning-enhanced super-resolution diffusion-weighted liver mri: improved image quality, diagnostic performance, and acceleration, Insights into Imaging 16 (2025).

[51] X. Li, Q. Liang, L. Zhuang, X. Zhang, T. Chen, L. Li, J. Liu, H. Calimente, Y. Wei, J. Hu, Preliminary study of mr diffusion tensor imaging of the liver for the diagnosis of hepatocellular carcinoma, PLoS ONE 10 (2015).

